# Crowdsourced EEG Experiments: A proof of concept for remote EEG acquisition using EmotivPRO Builder and EmotivLABS

**DOI:** 10.1101/2022.02.09.479644

**Authors:** Nikolas S Williams, William King, Geoffrey Mackellar, Roshini Randeniya, Alicia McCormick, Nicholas A Badcock

## Abstract

Online research platforms have enabled mass data collection enabling representative samples for cognitive behavioural studies. However, the benefits of online data collection have not been available for cognitive neuroscience fields such as electroencephalography (EEG). In this study, we introduce an approach for remote EEG data collection. We demonstrate how an experiment can be built via the EmotivPRO Builder and deployed to the EmotivLABS website where it can be completed by participants who own EMOTIV EEG headsets. To demonstrate the data collection technique, we collected EEG while participants engaged in a resting state task where participants sat with their eyes open and then eyes closed for two minutes each. We observed a significant difference in alpha power between the two conditions thereby demonstrating the well-known alpha suppression effect. Thus, we demonstrate that EEG data collection, particularly for frequency domain analysis, can be successfully conducted online with remote users.

## Introduction

The use of online research platforms for cognitive science has seen an increase in the past decade, with 10% to 30% of articles in cognitive science journals using online data collection marketplaces (Stewart et al., 2017). Currently, the most commonly used online participant recruitment agencies include Amazon Mechanical Turk (MTurk), Prolific Academic, and CrowdFlower. These platforms provide researchers access to between 10k - 500k of unique participants with a variety of demographic profiles (Peer et al., 2017). The use of these online marketplaces for cognitive science experiments has also been made more accessible to coding-naive researchers with javascript hosting platforms such as Pavlovia and Google SDK, and experiment builders such as PsychoPy3, jsPsych, and Gorilla Experiment Builder. Research has shown these platforms to be accurate and precise with regards to display duration and response time logging across many hardware configurations (Anwyl-Irvine et al., 2020). The proliferation of these online research tools has benefited researchers through access to larger and more diverse participant samples, increased experimental administration efficiency, and streamlined participant recruitment, all while yielding quality and reliable data similar to that collected in laboratory experiments (Finley & Penningroth, 2015; Germine et al., 2012).

A significant concern with in-person data collection is that the current literature is dominated by convenience samples consisting of narrow demographic profiles (Falk et al., 2013). Practical constraints of testing in traditional university laboratories often result in participant samples of convenience (e.g., first-year psychology students). Online testing alleviates some of this concern by giving access to a broader diversity of participants. Online testing also mitigates public health concerns as studies can be conducted with little to no physical contact. This is particularly important considering the COVID-19 pandemic in which experimental research was severely disrupted (Gentili & Cristea, 2020). By eliminating physical contact, online testing provides an alternative for continuing safe human-participant research during a public health crisis. While studies involving “hands-on” experiments, such as cognitive neuroscience research, have been disproportionately disrupted, unfortunately many of these types of experiments cannot be conducted online. This is because they use methods such as functional magnetic resonance imaging (fMRI), electroencephalography (EEG), and functional near-infrared spectroscopy (fNIRS) that require expensive, specialized devices that cannot be widely deployed or are unavailable to the general public.

Traditionally, EEG data is collected in a dedicated laboratory using expensive research-grade equipment that requires lengthy set up times, inordinate physical contact, and significant amounts of participant organization and administration. However, technological advances in the commercial EEG sector have resulted in the development of low-cost, wireless, consumer-grade EEG devices. Some of these systems have been adopted by the research community (see review by Sawangjai et al., 2020) resulting in more accessible and widespread EEG research. For example, Emotiv, Neurosky, InteraXon and OpenBCI have all released various consumer grade EEG systems that are all cost-effective (less than $1000 USD) and available to both scientific researchers as well as the general public.

EEG hardware advances only represent part of the solution with respect to online testing potential. Another requisite is an online platform capable of simultaneously presenting experiments and recording EEG data. Emotiv has recently developed an online platform, EmotivPRO Builder, that lets users build experiments within a browser-based graphical user interface (i.e., GUI). These experiments can then be published and made available to a contributor pool composed of Emotiv EEG system owners. Researchers can also use the EmotivLABS platform to collect data locally (for laboratory testing), making it a versatile tool for EEG experiment administration and data collection.

In the decade since the release of the first iteration, EPOC has been widely used by the scientific research community (for a review see Williams et al., 2020). EPOC is a 14-channel wireless EEG system that uses saline-soaked felt pads for signal conduction and has been empirically validated against research-grade systems (e.g., Neuroscan) and shown to record research-quality data (Badcock et al., 2013, 2015; de Lissa et al., 2015). Another system, EPOC Flex, is a 32-channel system that allows users to configure sensor placements within a traditional headcap. This system has also been validated against Neuroscan and shown to measure reliable auditory and visual event-related potentials (ERP), steady-state visual evoked potentials (SSVEP), and changes in alpha signatures (Williams, McArthur, de Wit, et al., 2020).

Taken together, the issues of representative sampling and public health clearly outline the benefits of an online, worldwide data collection platform that would increase the capacity to study underrepresented groups while also maintaining safe interactions during disease outbreaks. Thus, the purpose of this paper was to present a novel online EEG data collection platform that uses Emotiv EEG hardware and allows users to collect data remotely using the EmotivLABS platform. To do this, we used a simple resting state task to determine whether we could detect a well-known EEG phenomenon - alpha suppression - which is the decrease of alpha power when the eyes are open compared to when they are closed (Barry et al., 2007; Geller et al., 2014; Toscani et al., 2010).

## Method

### Participants

Participants were self-selected from the existing traffic on the EmotivLABS website. To contribute, participants must have possessed an Emotiv EEG headset. They also must have had an internet connection of minimum 5 Mbps. There was a one-off email communication sent to existing Emotiv headset owners in order to promote the study that included a link to the study. Participants were instructed to take the study at whatever time was suitable for them. The experiment was published to EmotivLABS on November 9th 2020 and remained available until August 14th 2021. Participants were not incentivised and participation was completely voluntary. This study was approved by the Ethics Committee of Macquarie University. All participants provided web-based informed consent before proceeding with the experiment.

### Data acquisition software: EmotivPRO Builder and EmotivLABS

The experiment was built using EmotivPRO Builder which is a web based interactive platform which allows users to build simple experiments using a graphical user interface. The finalized experiment was then published to EmotivLABS “Citizen Science” (https://labs.emotiv.com/), where the study was publicly visible to all visitors (see Figure 1). Thus, all instructions and stimuli were delivered to the participant through this online platform.

**Figure 1:**
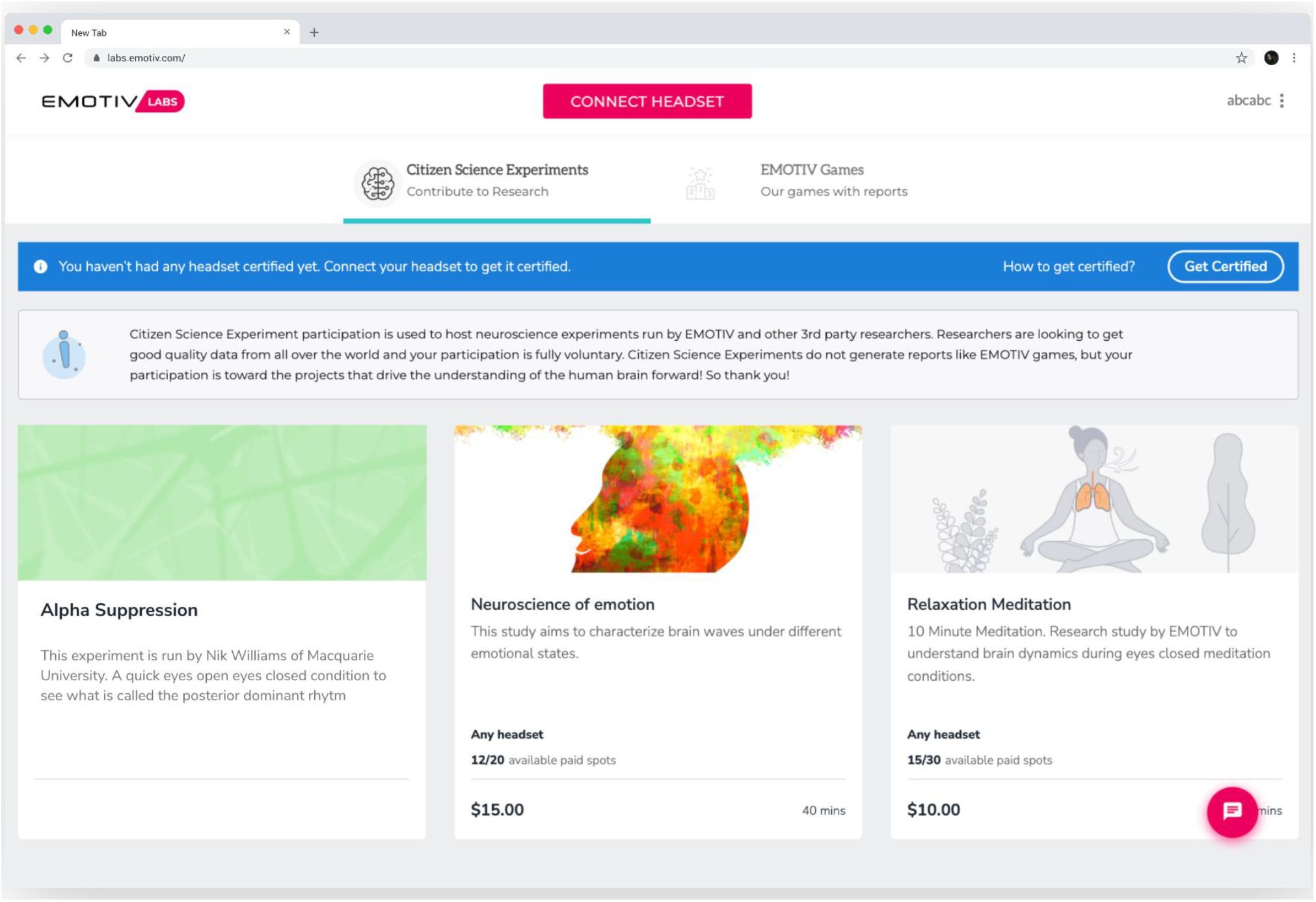
EmotivLABS Citizen Science platform. Participants can log on to the EmotivLABS Citizen Science platform using their EmotivID and participate in the experiments shown.

### Data acquisition devices: EEG devices

Participants in this study used one of the following EEG devices: EMOTIV INSIGHT, EPOC, EPOC+, or EPOCX. Although EPOC, EPOC+, and EPOCX are different models, they have a similar form factor and sensor configuration and only differ slightly in their technical specifications. Thus, for simplicity we subsequently refer to all three systems collectively as EPOC.

All Emotiv systems are wireless and connect to the participant’s computer through EmotivPRO software using either an Emotiv USB Receiver Dongle or the computer’s native Bluetooth adapter (See **Table 1** for headset details). The EPOC and INSIGHT headsets sample the EEG data internally at 2048KHz. Before the data is transmitted wirelessly to the recording computer, the data is downsampled to either 256Hz or 128Hz. This is done after a dual notch filter at 50Hz and 60Hz and an antialiasing filter has been applied to reduce mains-line noise and the aliasing of frequencies higher than 64Hz into the transmitted data.

**Table 1:**
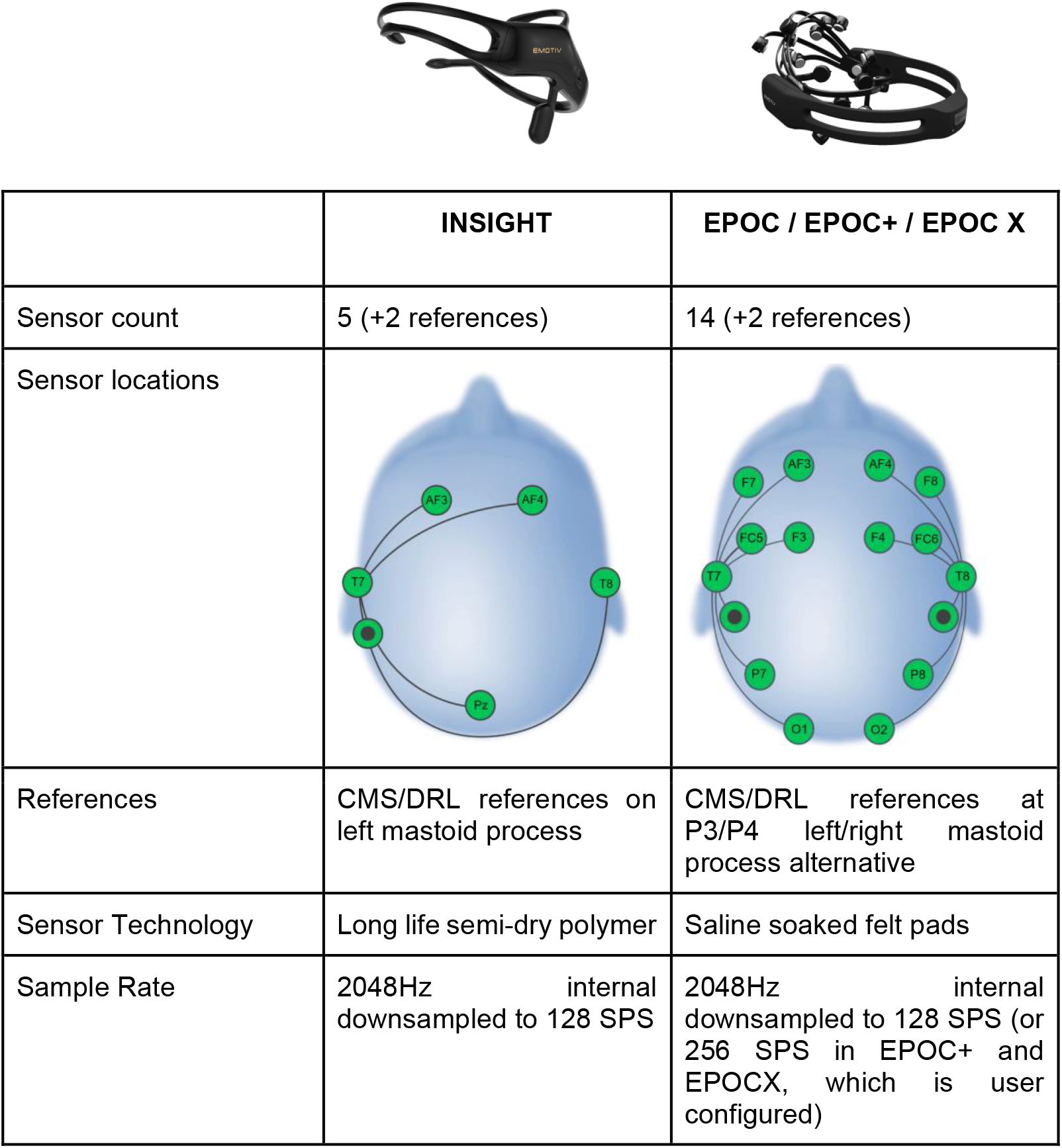
EEG devices used in the study.

|  | INSIGHT | EPOC / EPOC+ / EPOC X |
| --- | --- | --- |
| Sensor count | 5 (+2 references) | 14 (+2 references) |
| Sensor locations |  |  |
| References | CMS/DRL references on left mastoid process | CMS/DRL references at P3/P4 left/right mastoid process alternative |
| Sensor Technology | Long life semi-dry polymer | Saline soaked felt pads |
| Sample Rate | 2048Hz internal downsampled to 128 SPS | 2048Hz internal downsampled to 128 SPS (or 256 SPS in EPOC+ and EPOCX, which is user configured) |

### Data acquisition: Procedure

After initiating the experiment on EmotivLABS, the participant was guided through the headset connection process. To ensure data quality, the participant is required to pass an EEG quality (EQ) “gate”.

EQ is a gaussian algorithm based on EEG signal characteristics, contact quality (CQ), and EEG sample loss rate. CQ is a direct measure of the resistance of the circuit through each electrode and the driven right leg (DRL) electrode. For efficient removal of the common mode noise, we also need to ensure that the common mode signal (CMS) electrode has good electrical contact with the skin. The EQ model was built on the analysis of approximately 40 records of 20-40 minutes in which bad and good EEG segments were visually inspected and labeled. The algorithm uses the model to rank a block of two-second EEG signals as very good, good, poor or very bad (Figure 2). To pass the EQ gate, participants must exceed 82%. Please note that high EQ does not guarantee artifact-free data.

**Figure 2:**
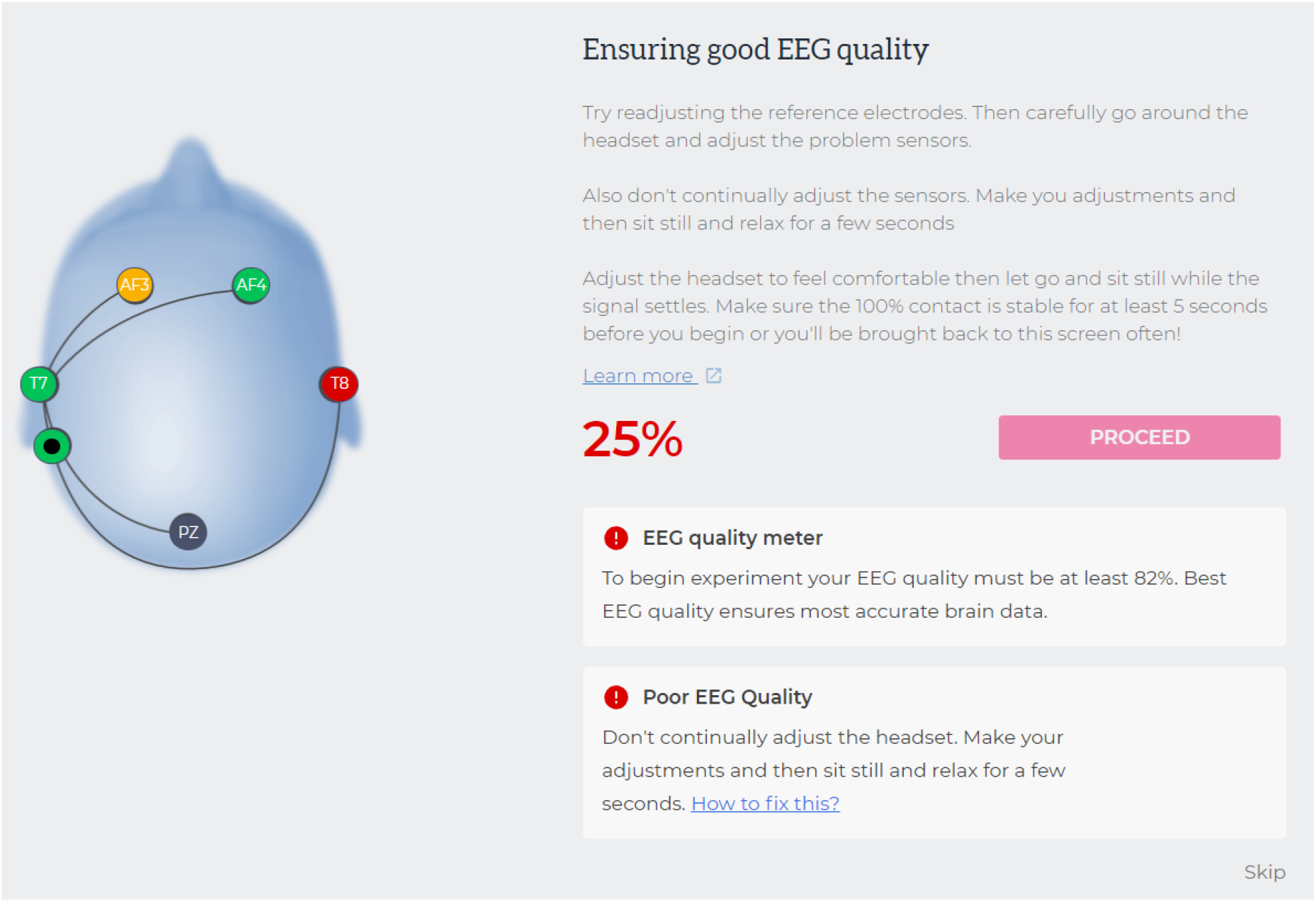
Participants are required to pass an EEG quality gate. Participants see an interactive headmap (left) with a colour indicator of EEG quality (black - bad, red - poor, orange - good, green - very good) at each sensor and must achieve more than 80% overall quality to be included in the study.

After passing the EEG Quality Gate, participants were presented with a digital copy of the information and consent form. Participants gave consent by ticking a “yes I provide consent” box and clicking the “next” button.

Participants were next encouraged to ensure they will not be disturbed, to turn off their phone (or place in airplane mode), to check that their computer and EEG headset are adequately charged, and that their computer sound is turned up. The participant was also asked to switch their browser to full screen mode to minimize any disturbances from other computer applications.

Participants then began the experiment and were presented with a three-second audio-visual countdown after which a fixation cross appeared in the middle of the screen. Participants were instructed to keep their eyes open during this time. The Eyes Open condition lasted two minutes. Next, participants were instructed to close their eyes until they heard a tone, which signaled the end of the trial. The Eyes Closed condition also lasted two minutes. The conditions were not counter-balanced and participants always completed Eyes Open first.

### Data Processing and Analysis

Participant data were first downsampled to 128 Hz if needed and re-referenced to the interquartile mean of all the channels. The data were then high-pass filtered at a threshold of 0.5 Hz. The absolute power (in dB scale) for each two-minute epoch (Eyes Open/Eyes Closed) was calculated using Fast Fourier Transform (FFT) for all electrodes in the alpha band (8-12Hz). Custom scripts in python were used for EEG processing and FFT and have been made openly available (find link to repository under Data Availability).

We performed a paired samples t-test and Bayesian t-test on the absolute Alpha power between the Eyes Open and Eyes Closed conditions measured at Pz, O1, and O2. We chose these sites as visual alpha suppression is typically larger at posterior sites (Magosso et al., 2019; Thut et al., 2006; Toscani et al., 2010). To determine the strength of evidence provided by the Bayes Factor calculated in the t-test, we used the guidelines reported by (Jarosz & Wiley, 2014). We also report mean alpha averaged over all electrodes for INSIGHT and EPOC separately. Statistical analysis was undertaken using R (R Core Team, 2021) and all figures are presented using ggplot (Wickham, 2016).

## Results

### Sample size

During the period in which the experiment was active on EmotivLABS website, 105 recordings were initiated by 60 unique participants. Of the 60 unique participants, only 28 participants successfully completed both Eyes Open and Eyes Closed conditions. For cases in which a single participant made two successful recordings, the first recording was retained and the second was excluded in the analysis. Overall, we received data from four INSIGHT, one EPOC, six EPOC+, and six EPOCX systems. We also received data from one EPOCFLEX, but it was excluded from analysis as it cannot be pooled with the other headsets due to the differing sensor count. Thus, 27 complete EEG recordings from 27 unique participants were processed and analyzed. Statistical analysis was conducted separately for INSIGHT (N = 14) and EPOC/EPOC+/EPOCX (N = 13) headsets.

Overall, there were 77 recordings that did not meet the criteria for this study. In 14 cases participants skipped the EEG quality gate. Thirty-seven recordings were abandoned without completing the EQ gate. In another 26 recordings participants passed the gate but abandoned the experiment at various stages. Finally, one recording was a duplicate where a participant completed the experiment twice.

### Participant demographics

Of the 28 participants, 26 specified their age. These ranged from 18 to 64 (M = 37.8, SD = 12.6). Genders included 22 males, 5 females, and one unspecified gender. Twenty-three participants were right-handed, four were left-handed, and one ambidextrous based on self-reporting writing hand. Education levels ranged from high school to doctoral degree (see Figure 3 and Table 2 for demographic profiles). Participant recordings were initiated from the United States (n = 8), Australia (n = 3), Sweden (n = 2), Argentina (n = 2), Iran (n = 2) and one each from Czech Republic, Egypt, Spain, United Kingdom, Singapore, Turkey, Lithuania, China, Finland, Mexico, and an unspecified country.

**Table 2:** Demographic profile of participants.

| Factor | N | % of total sample (N = 27) |
| --- | --- | --- |
| **Handedness** |  |  |
| - Right | 23 | 82.14 |
| - Left | 4 | 14.29 |
| - Ambidextrous | 1 | 3.57 |
| **Education Level** |  |  |
| - Below High School | 1 | 3.57 |
| - High school or equivalent | 4 | 14.29 |
| - Bachelor's degree | 9 | 32.14 |
| - Vocational/technical training | 5 | 17.86 |
| - Professional degree | 1 | 3.57 |
| - Masters degree | 5 | 17.86 |
| - Doctoral degree | 3 | 10.71 |

**Figure 3:**
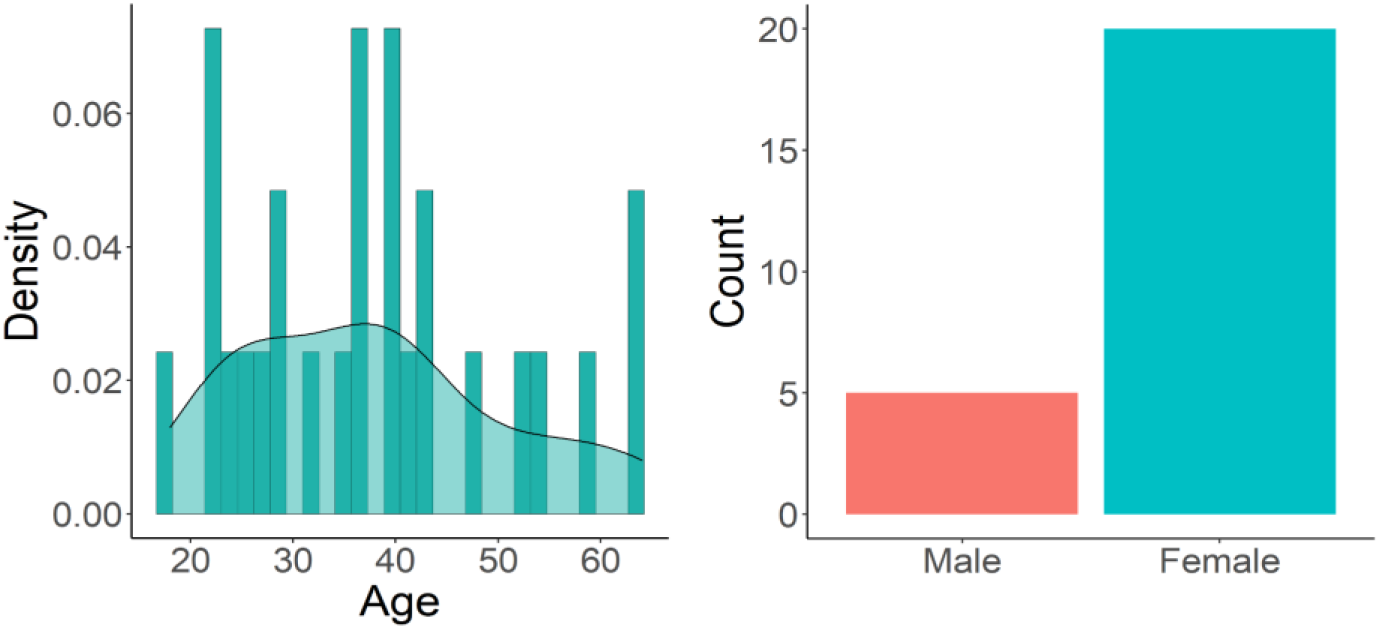
Demographic profile of participants.

### Alpha Suppression

We observed strong evidence for a difference in mean alpha power between Eyes Open and Eyes Closed conditions for INSIGHT (*t*(13) = 4.78, *p* = < .001, Cohen’s *d* = 1.28, BF_10_ = 94.27) and for EPOC (*t*(11) = 4.952, *p* = < .001, Cohen’s *d* = 1.43, BF_10_ = 81.50). See Figure 4 for data distributions and descriptive statistics. Also see Figure 5 for topographic alpha power distributions.

**Figure 4:**
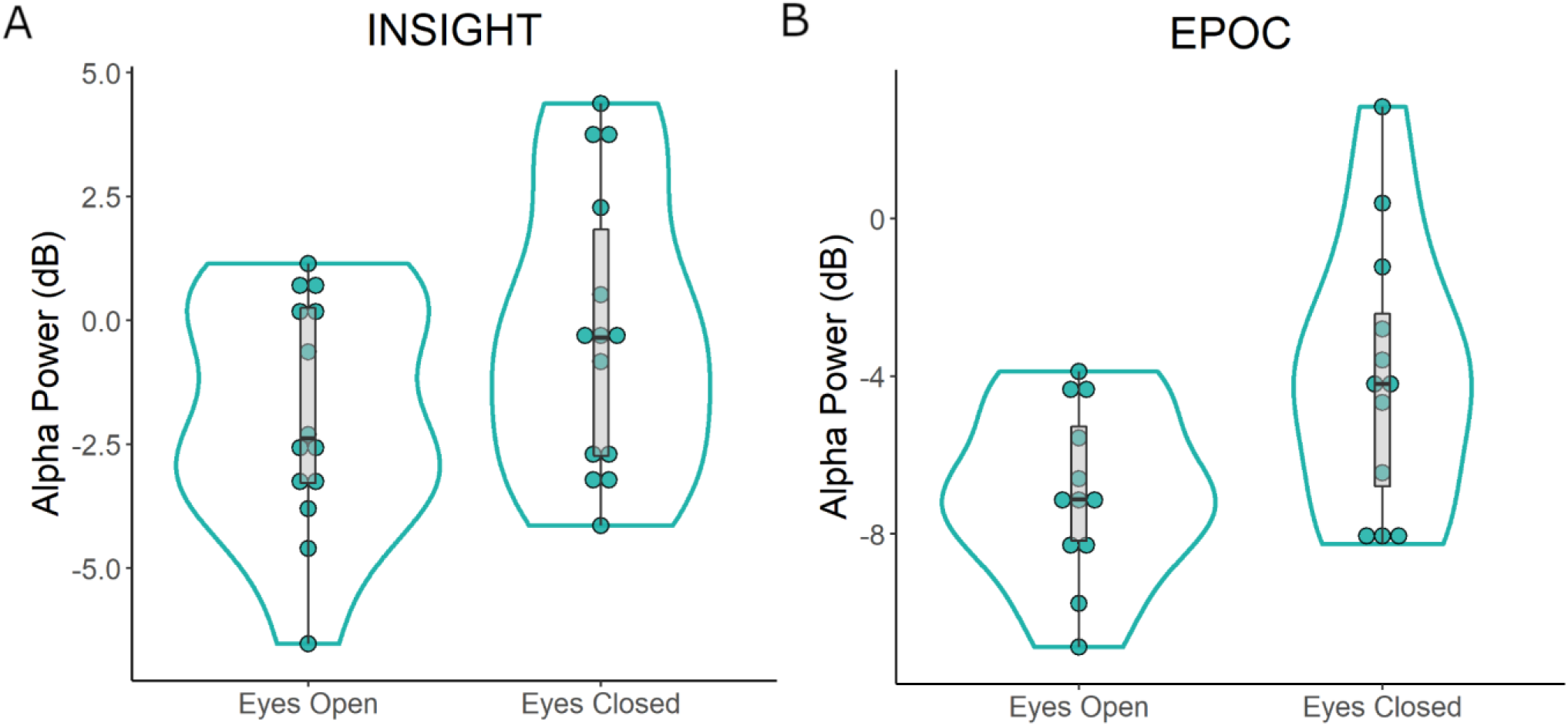
Alpha suppression (between Eyes Open and Eyes Closed) conditions can be observed in the global mean (i.e., alpha power averaged over all electrodes) for A) INSIGHT and B) EPOC headsets. Violin plots show a density curve (blue outline) with each dot representing an individual participant. Boxplots show median and interquartile ranges and min and max values.

**Figure 5.**
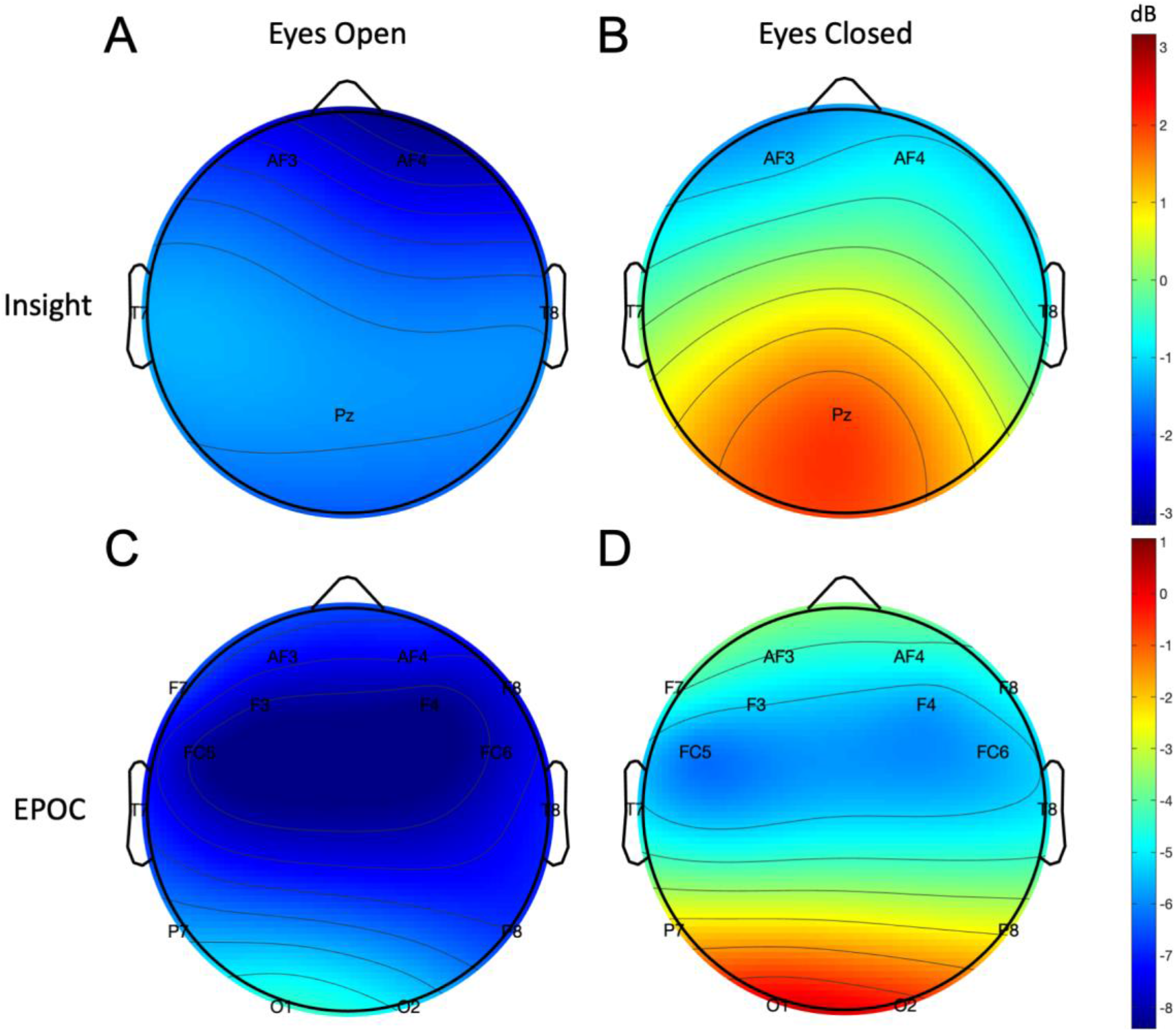
Topographical alpha power distributions measured by Insight (A and B) and EPOC (C and D) in the Eyes Open (left column) and Eyes Closed (right column) conditions.

We also observed strong evidence for difference in alpha power between Eyes Open and Eyes Closed conditions at Pz electrode for INSIGHT (*t*(13) = 4.95, *p* < .001, *d* = 1.32, BF_10_ = 123.29) and at O1 (*t*(11) = 4.52, *p* < .001, *d* = 1.31, BF_10_ = 45.34) and O2 (*t*(11) = 4.74, *p* = < .001, *d* = 1.37, BF_10_ = 60.78) for EPOC. See Figure 6 for data distributions and descriptive statistics.

**Figure 6:**
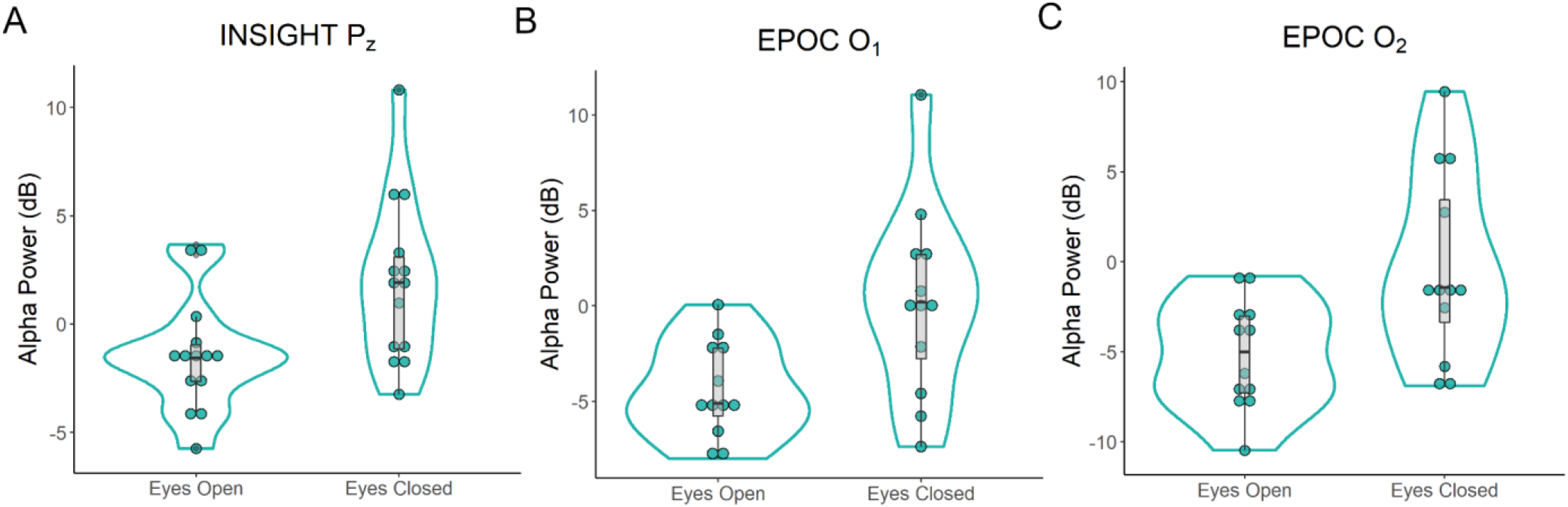
Alpha suppression (between Eyes Open and Eyes Closed) conditions can be observed at posterior electrodes Pz of INSIGHT(A), O1 of EPOC (B), and O2 of EPOC (C).

## Discussion

In this study, we introduced an approach for remote EEG data collection. We demonstrated how an experiment can be built via the EmotivPRO Builder and deployed to home users that own EEG headsets to undertake experiments. We demonstrated alpha suppression phenomenon with home users as previously observed in laboratory settings (Bacigalupo & Luck, 2019; Magosso et al., 2019; Toscani et al., 2010), thus demonstrating the feasibility of conducting online studies to collect EEG data.

The method described for collecting online EEG data has several strengths in ensuring participants obtain high quality EEG data. The EEG quality gate ensures that participants achieve high contact quality (low impedance) at each channel and enables participants to identify and improve signal quality before they can proceed. However, we note that while 60 unique participants initiated this study on the EmotivLABS website, only 28 completed it. As noted in the results, some participants were unable to pass the EQ gate and some participants abandoned the experiment midway. Home users may require additional training on obtaining high quality EEG data in order to minimize time spent at the EQ gate thereby motivating them to proceed with an experiment. Additionally, since this was a voluntary study, providing incentives (e.g., monetary compensation) may motivate participants to complete the EQ gate and complete experiments.

A particular benefit of online studies is the ability to sample a wider range of demographics (Stewart et al., 2017) than is typically possible with laboratory EEG systems. Our study was able to reach a variety of participants from geographical locations (European, Asian, Australian, and American), ranging ages 18 to 64 and consisting of educational backgrounds arising from high school to professional degrees. However, the gender ratio in our sample was biased towards males. While our sample size was small, there is potential for low-cost EEG systems to proliferate thereby increasing the participant pool available to users of EmotivLABS. This would result in further sample diversity and shortened data collection times.

The method introduced in this study showed the feasibility of conducting resting state tasks and frequency domain analysis. We did not investigate event-related potential (ERP) experiments in this study which require accurate timestamping to enable ERP component analysis. However, it is currently possible to integrate PsychoPy with EmotivPRO Builder to deliver ERP stimuli and obtain accurate timestamps. Future online ERP studies should be conducted to determine the reliability and accuracy of such data.

Our feasibility study required very simple tasks of the participant with clear instructions. We demonstrated that participants can undertake simple studies. However, a limitation for conducting online EEG data collection is that of conducting complex tasks. It has been demonstrated that as task complexity increases, participants have difficulty in understanding task instructions online compared to lab as they do not have the opportunity to clarify (Finley & Penningroth, 2015). This is a difficulty relevant to all online data collection platforms but may be overcome with significant pilot testing online, revising of experiment instructions and guidelines, as well as using video-conferencing to provide specific instructions and guidance to participants.

While the online platform gives as much direction as possible for placement of electrodes, there may still be differences in where sensors are placed on the participant’s head. In laboratory settings this is avoided by having trained researchers conduct the EEG setup. However, more instruction and training may be required for home users. In addition, head sizes differ. Whereas this is typically catered for in a lab environment by having different sized caps available, EMOTIV systems are available in a single size only. As such, they may not always conform to maintain exact 10-20 placement over the complete range of head sizes and shapes. Thus, sensor placement may vary by a small degree from participant to participant and we could not ensure precise sensor placement. However, the EMOTIV systems have flexible arms that have been designed to accommodate the heads of participants from small adolescents to large adults. Thus, any deviances from exact 10-20 specifications were likely to be very small.

In this paper, we demonstrate the potential for obtaining high quality EEG data via an online platform. The benefits of such testing are notable when considering current concerns with sampling diversity and public health. By leveraging a user base that is familiar with EEG systems and online platforms, researchers may be able to conduct global EEG studies in a manner that has not been possible.

## Data availability

Raw EEG data for all 27 participants and preprocessing, analysis and visualization scripts have been made openly available via OSF here: https://osf.io/9bvgh/?view_only=70744f62157c46d5bd731480db1873df

## References

R Core Team (2021). R: A language and environment for statistical computing. R Foundation for Statistical Computing, Vienna, Austria. URL https://www.R-project.org/ https://doi.org/10.1007/s00221-010-2444-7

Wickham, H. (2016). ggplot2: Elegant Graphics for Data Analysis. Springer-Verlag New York. https://ggplot2.tidyverse.org.

Anwyl-Irvine, A., Dalmaijer, E. S., Hodges, N., & Evershed, J. K. (2020). Realistic precision and accuracy of online experiment platforms, web browsers, and devices. Behavior Research Methods. https://doi.org/10.3758/s13428-020-01501-5

Bacigalupo, F., & Luck, S. J. (2019). Lateralized Suppression of Alpha-Band EEG Activity As a Mechanism of Target Processing. The Journal of Neuroscience, 39(5), 900–917. https://doi.org/10.1523/JNEUROSCI.0183-18.2018

Badcock, N. A., Mousikou, P., Mahajan, Y., de Lissa, P., Thie, J., & McArthur, G. (2013). Validation of the Emotiv EPOC® EEG gaming system for measuring research quality auditory ERPs. PeerJ, 1, e38. https://doi.org/10.7717/peerj.38

Badcock, N. A., Preece, K. A. Wit, B. de, Glenn, K., Fieder, N., Thie, J., & McArthur, G. (2015). Validation of the Emotiv EPOC EEG system for research quality auditory event-related potentials in children. PeerJ, 3, e907. https://doi.org/10.7717/peerj.907

Barry, R. J., Clarke, A. R., Johnstone, S. J., Magee, C. A., & Rushby, J. A. (2007). EEG differences between eyes-closed and eyes-open resting conditions. Clinical Neurophysiology: Official Journal of the International Federation of Clinical Neurophysiology, 118(12), 2765–2773. https://doi.org/10.1016/j.clinph.2007.07.028

de Lissa, P., Sörensen, S., Badcock, N., Thie, J., & McArthur, G. (2015). Measuring the face-sensitive N170 with a gaming EEG system: A validation study. Journal of Neuroscience Methods, 253, 47–54. https://doi.org/10.1016/j.jneumeth.2015.05.025

Falk, E. B., Hyde, L. W., Mitchell, C., Faul, J., Gonzalez, R., Heitzeg, M. M., Keating, D. P., Langa, K. M., Martz, M. E., Maslowsky, J., Morrison, F. J., Noll, D. C., Patrick, M. E., Pfeffer, F. T., Reuter-Lorenz, P. A., Thomason, M. E., Davis-Kean, P., Monk, C. S., & Schulenberg, J. (2013). What is a representative brain? Neuroscience meets population science. Proceedings of the National Academy of Sciences, 110(44), 17615–17622. https://doi.org/10.1073/pnas.1310134110

Finley, A., & Penningroth, S. (2015). Online versus In-lab: Pros and Cons of an Online Prospective Memory Experiment (pp. 135–161).

Geller, A. S., Burke, J. F., Sperling, M. R., Sharan, A. D., Litt, B., Baltuch, G. H., Lucas, T. H., & Kahana, M. J. (2014). Eye closure causes widespread low-frequency power increase and focal gamma attenuation in the human electrocorticogram. Clinical Neurophysiology: Official Journal of the International Federation of Clinical Neurophysiology, 125(9), 1764–1773. https://doi.org/10.1016/j.clinph.2014.01.021

Gentili, C., & Cristea, I. A. (2020). Challenges and Opportunities for Human Behavior Research in the Coronavirus Disease (COVID-19) Pandemic. Frontiers in Psychology, 11, 1786. https://doi.org/10.3389/fpsyg.2020.01786

Germine, L., Nakayama, K., Duchaine, B. C., Chabris, C. F., Chatterjee, G., & Wilmer, J. B. (2012). Is the Web as good as the lab? Comparable performance from Web and lab in cognitive/perceptual experiments. Psychonomic Bulletin & Review, 19(5), 847–857. https://doi.org/10.3758/s13423-012-0296-9

Jarosz, A., & Wiley, J. (2014). What Are the Odds? A Practical Guide to Computing and Reporting Bayes Factors. The Journal of Problem Solving, 7(1). https://doi.org/10.7771/1932-6246.1167

Magosso, E., De Crescenzio, F., Ricci, G., Piastra, S., & Ursino, M. (2019). EEG Alpha Power Is Modulated by Attentional Changes during Cognitive Tasks and Virtual Reality Immersion. Computational Intelligence and Neuroscience, 2019, e7051079. https://doi.org/10.1155/2019/7051079

Peer, E., Brandimarte, L., Samat, S., & Acquisti, A. (2017). Beyond the Turk: Alternative platforms for crowdsourcing behavioral research. Journal of Experimental Social Psychology, 70, 153–163. https://doi.org/10.1016/j.jesp.2017.01.006

R Core Team. (2021). R: A language and environment for statistical computing. R Foundation for Statistical Computing. https://www.R-project.org/

Sawangjai, P., Hompoonsup, S., Leelaarporn, P., Kongwudhikunakorn, S., & Wilaiprasitporn, T. (2020). Consumer Grade EEG Measuring Sensors as Research Tools: A Review. IEEE Sensors Journal, 20(8), 3996–4024. https://doi.org/10.1109/JSEN.2019.2962874

Stewart, N., Chandler, J., & Paolacci, G. (2017). Crowdsourcing Samples in Cognitive Science. Trends in Cognitive Sciences, 21(10), 736–748. https://doi.org/10.1016/j.tics.2017.06.007

Thut, G., Nietzel, A., Brandt, S. A., & Pascual-Leone, A. (2006). α-Band Electroencephalographic Activity over Occipital Cortex Indexes Visuospatial Attention Bias and Predicts Visual Target Detection. Journal of Neuroscience, 26(37), 9494–9502. https://doi.org/10.1523/JNEUROSCI.0875-06.2006

Toscani, M., Marzi, T., Righi, S., Viggiano, M. P., & Baldassi, S. (2010). Alpha waves: A neural signature of visual suppression. Experimental Brain Research, 207(3–4), 213–219. https://doi.org/10.1007/s00221-010-2444-7

Wickham, H. (2016). ggplot2: Elegant Graphics for Data Analysis. Springer-Verlag. https://ggplot2.tidyverse.org

Williams, N. S., McArthur, G. M., & Badcock, N. A. (2020). 10 years of EPOC: A scoping review of Emotiv’s portable EEG device. BioRxiv, 2020.07.14.202085. https://doi.org/10.1101/2020.07.14.202085

Williams, N. S., McArthur, G. M., de Wit, B., Ibrahim, G., & Badcock, N. A. (2020). A validation of Emotiv EPOC Flex saline for EEG and ERP research. PeerJ, 8, e9713. https://doi.org/10.7717/peerj.9713

